## Supplementary Information for "Structure of the stress-related LHCSR1 complex determined by an integrated computational strategy"

#### Supplementary Methods

**Spectral densities** The spectral density of the Chl excited states was modeled as a sum of overdamped Brownian oscillator and 48 high-frequency modes. The parameters for the spectral density were taken from the fitting of the fluorescence line narrowing experimental data for LHCII by Novoderezhkin et al.[1]. The spectral density of the Car $S_2$ excited state has been modeled by a sum of Brownian oscillator and two high frequency modes. The high frequency modes of the Cars were described by underdamped Brownian oscillator spectral density with the two most pronounced vibrational modes at $1050 cm^{-1}$ and $1600 cm^{-1}$, which corresponds to C-C and C=C stretch frequencies of the $\pi$-conjugated chain which are coupled to the optical transition[2]. The Huang-Rhys factors and corresponding reorganization energies $\lambda$ of the individual vibrations and parameters of the Brownian oscillator were obtained from fitting the optical spectra of the isolated carotenoids in n-hexane as $\lambda_{1050} = 400cm^{-1}$ and $\lambda_{1600} = 1600cm^{-1}$. The low frequency modes were modeled with the overdamped Brownian oscillator spectral density with correlation time $\tau_c = 200fs$ and reorganization energy $\lambda = 250cm^{-1}$. Parameters of the low frequency part of the spectral density were obtained from fitting of the width of the experimental spectra. The comparison of the Car spectra from the fitted spectral density and the experimental one can be found in Figure S20.

**Site energies and static disorder** For obtaining the pigment site energies we used a two-step approach, computing the excitation energies for the pigments in the protein environment and *in vacuo*, so as to separate the geometrical and environment contributions, as in our previous work[3]. The environment contribution is obtained as the difference between *in protein* and *in vacuo* excitation energies. The geometrical contribution arises from the distortion of bond lengths, angles, and dihedrals with respect to an optimized Chl structure. As previously found, a large variability in the excitation energy comes from the bond lengths[4]. As the bond length variability is connected with "fast" degrees of freedom, which are included in the spectral density, we adopted a statistical protocol to remove the bond-length contribution ($\Delta E^{\text{bond}}$) from the excitation energies. The bond-length contribution was estimated through a multilinear fitting, as done in ref. 4, and then averaged over all chlorophylls for all investigated frames. For each pigment in each frame, the bond-length contribution estimated from the fitting was subtracted from the excitation energy. The remaining geometrical contribution (distortion) was averaged only over the structures of each pigment in each cluster because the average geometry could be statistically different in different pigments and conformations. The environmental contribution was computed for every single frame separately, which leads to correct thermal sampling of the excitation energy disorder. With this scheme, we (i) correct the intrinsic error of the MD simulation for the pigment bond lengths, (ii) exclude double-counting of bond length contribution to the energy disorder, and (iii) reduce the "noise" in the excitation energies coming only from the limited sampling of pigment geometries. The final corrected site energy for each pigment $j$ in each configuration $t$ reads:

$$E_{jt}^{\text{corrected}} = E_0 + \langle \Delta E^{\text{bond}} \rangle + \langle \Delta E_j^{\text{distortion}} \rangle + \Delta E_{jt}^{\text{env}} \quad \text{(S1)}$$

where $E_0$ is the excitation energy of a reference pigment structure, $\langle \Delta E^{\text{bond}} \rangle$ is the average bond-length contribution, $\langle \Delta E_j^{\text{distortion}} \rangle$ is the pigment-specific average distortion contribution of pigment $j$, and $\Delta E_{jt}^{\text{env}}$ is the environment contribution for pigment $j$ in configuration $t$. In this scheme, the Hamiltonian disorder is sampled directly from the distribution of $\Delta E^{\text{env}}$ rather than extracted from a predefined distribution.

The linear regression using all bond lengths explained more than 80% (computed as: $1 - \langle (E_{\text{bond}} - E)^2 \rangle / \langle (E - \bar{E})^2 \rangle$) of the variability of the Chls excitation energies *in vacuo*, Figure S18. As in classical force fields, the bond lengths should sample the same distribution, we expect the bond length contribution to average out. However, since this contribution has a major impact on the site energy fluctuations, it obscures the variability arising from the environment's conformation. Based on the linear regression, however, we were able to subtract the bond length contributions, and thus remove the large uncertainty in excitation energies coming from the fluctuations of bond lengths, Figure S19. Site energies corrected according to Eq. S1, shown in Figure S15, were used to build the disordered exciton Hamiltonian for each cluster.

We note that the site energies of the whole chlorophyll system were moved by $-1381$ cm$^{-1}$ in order to account for the systematic error of the DFT approach and MD pigment geometries. The value of the site energy shift was independently determined in our previous study of the LHCII[3] and corrected to the smaller basis set used in this work to ensure the consistency between the studied light-harvesting antennas. The basis set correction was determined by comparison of the site energies of several selected chlorophylls computed for two replicas (100 frames) with M06-2X functional and 6-31+G(d) basis set, as used in our previous study of the LHCII[3], and the 6-31G(d) basis used in the current work. The site energies for the M06-2X/6-31+G(d) were systematically lower by 312 cm$^{-1}$ with respect to the 6-31G(d) basis set. The site energy of carotenoids was moved by 1583 cm$^{-1}$ for the same reason as chlorophylls (Table S1). This value for the shift was determined by comparison of the experimental and QM energy gap between chlorophyll $Q_y$ transition and carotenoid $S_2$ transition for isolated pigments in n-hexane. For CP29, Chl b transition energies were shifted by $+200$cm$^{-1}$ to correct the intrinsic error of the TD-DFT method in estimation of the energy gap between Chl a and Chl b (See Supplementary Methods).

#### Supplementary Figures

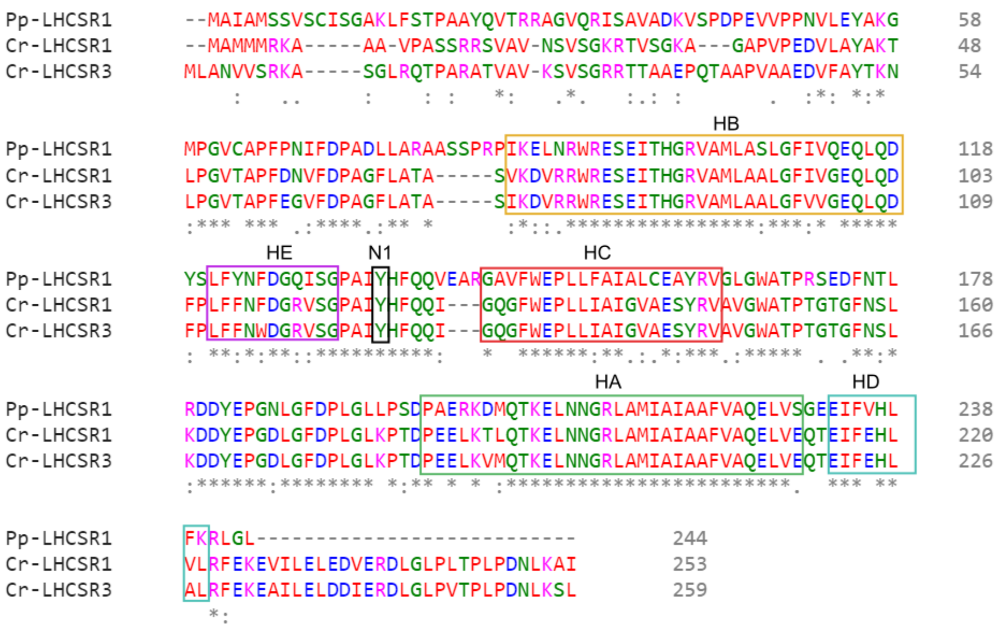


Figure S1: Alignment of LHCSR sequences for LHCSR1 from *P. Patens* (Pp-LHCSR1) and LHCSR1 and LHCSR3 from *C. Reinhardtii* (Cr-LHCSR1, Cr-LHCSR3). Approximate positions of helices A-E are indicated by the square boxes. N1 indicates the conserved Tyr that constitutes the putative binding site for N1-Lut.

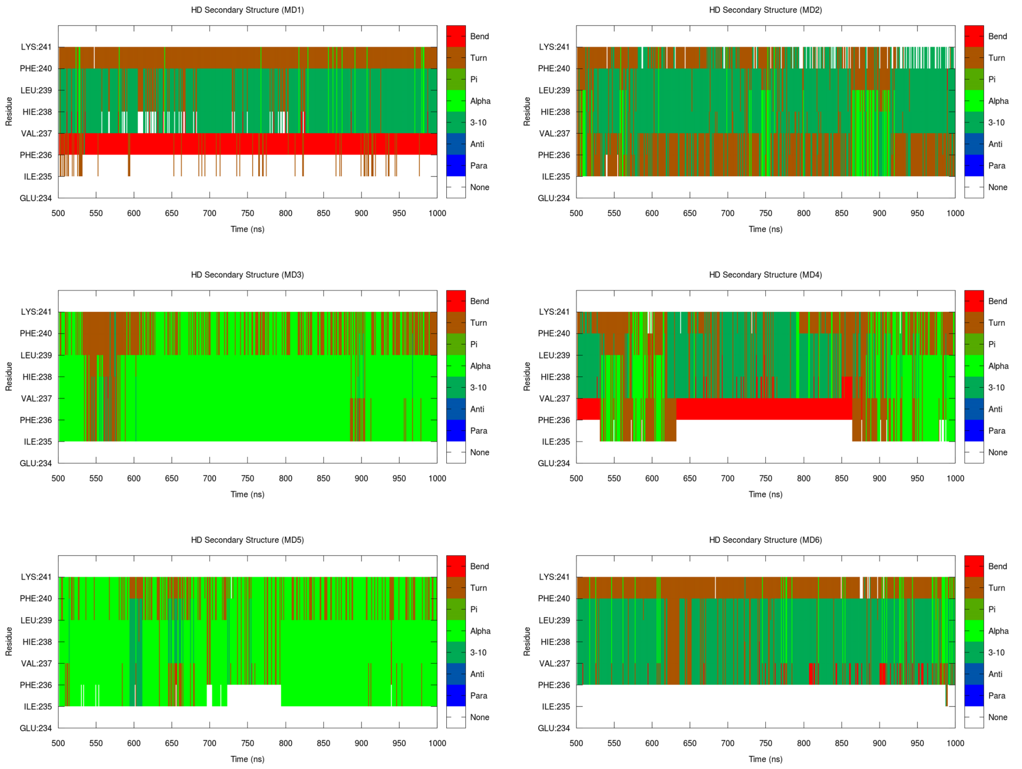


Figure S2: Secondary structure in the Helix D region on refinement MDs 1-6.

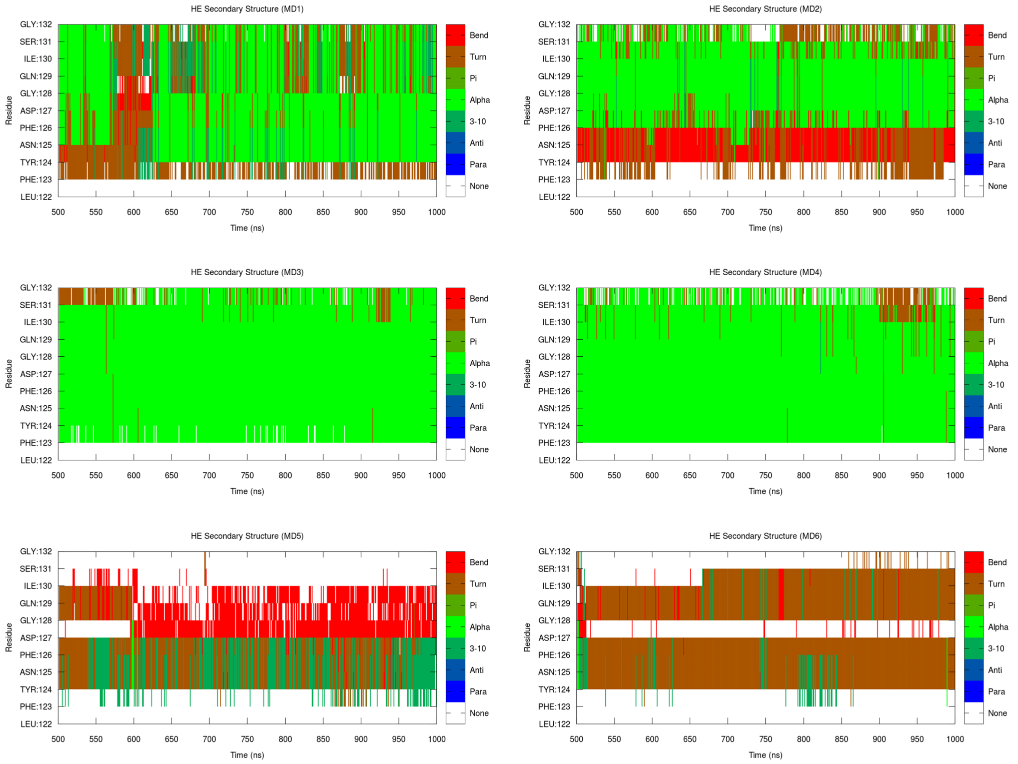


Figure S3: Secondary structure in the Helix E region on refinement MDs 1-6.

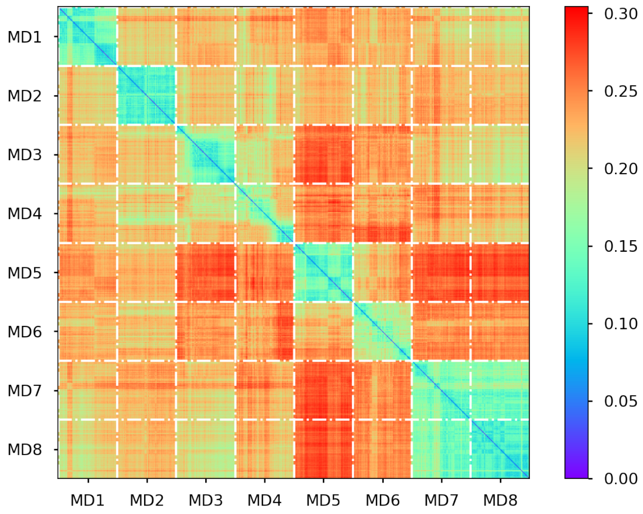


Figure S4: RMSD in the Ramachandran torsional space between frames of the six replicas. 100 structures from the last 500 ns of each replica were included in this RMSD matrix. White lines separate the different replicas. Blue-green regions indicate pairs of structures that share a greater similarity.

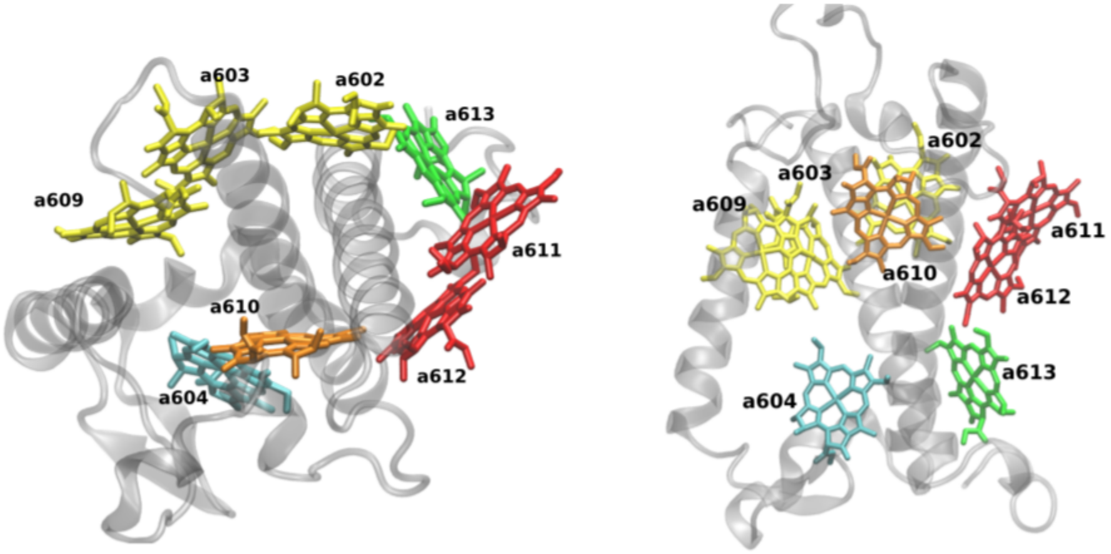


Figure S5: Visualization of the LHCSR1 structure with the chlorophyll names

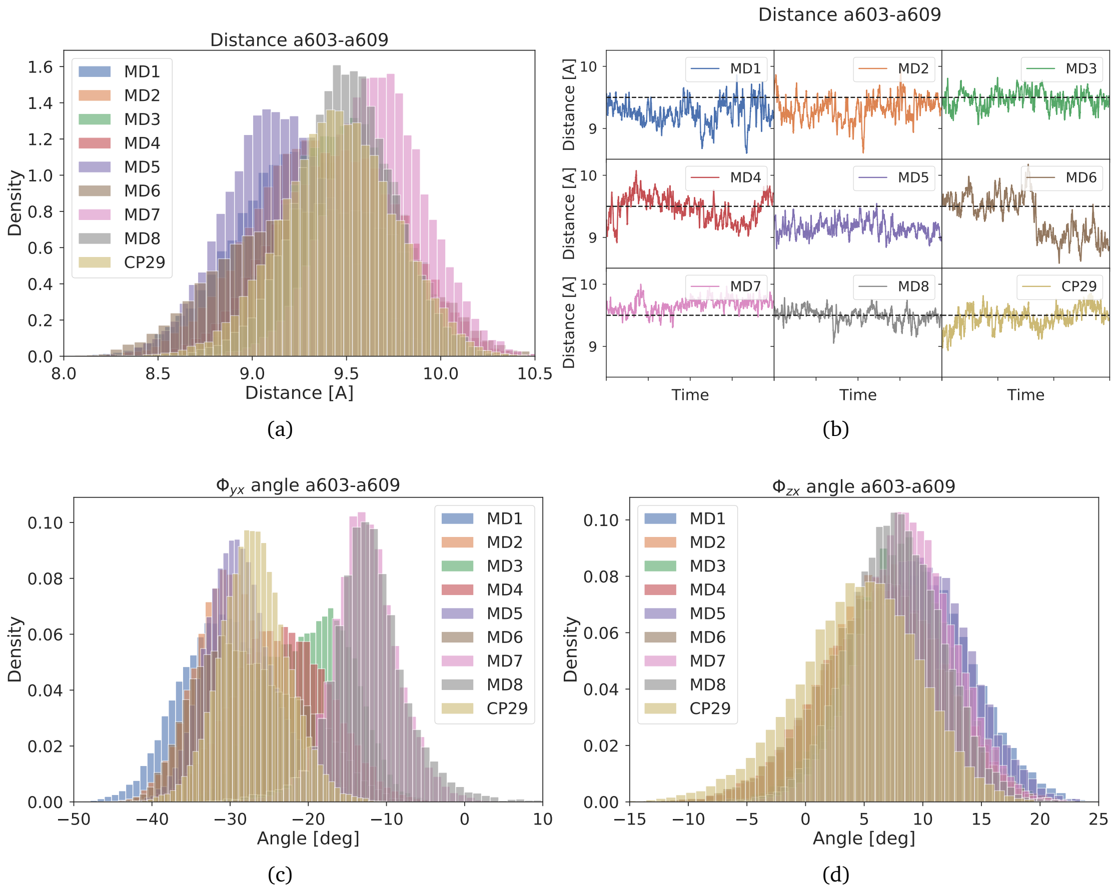


Figure S6: a-b) Mutual distance between central Mg site of the chlorophyll a603 and a609 for the individual replicas (MD) of the LHCSR1 and CP29. For the time evolution, we plotted running average over 100 frames c) Mutual rotation of the chlorophylls in the plane of the chlorophyll conjugated ring. d) Mutual rotation of the chlorophylls in the plane perpendicular to the conjugated ring.

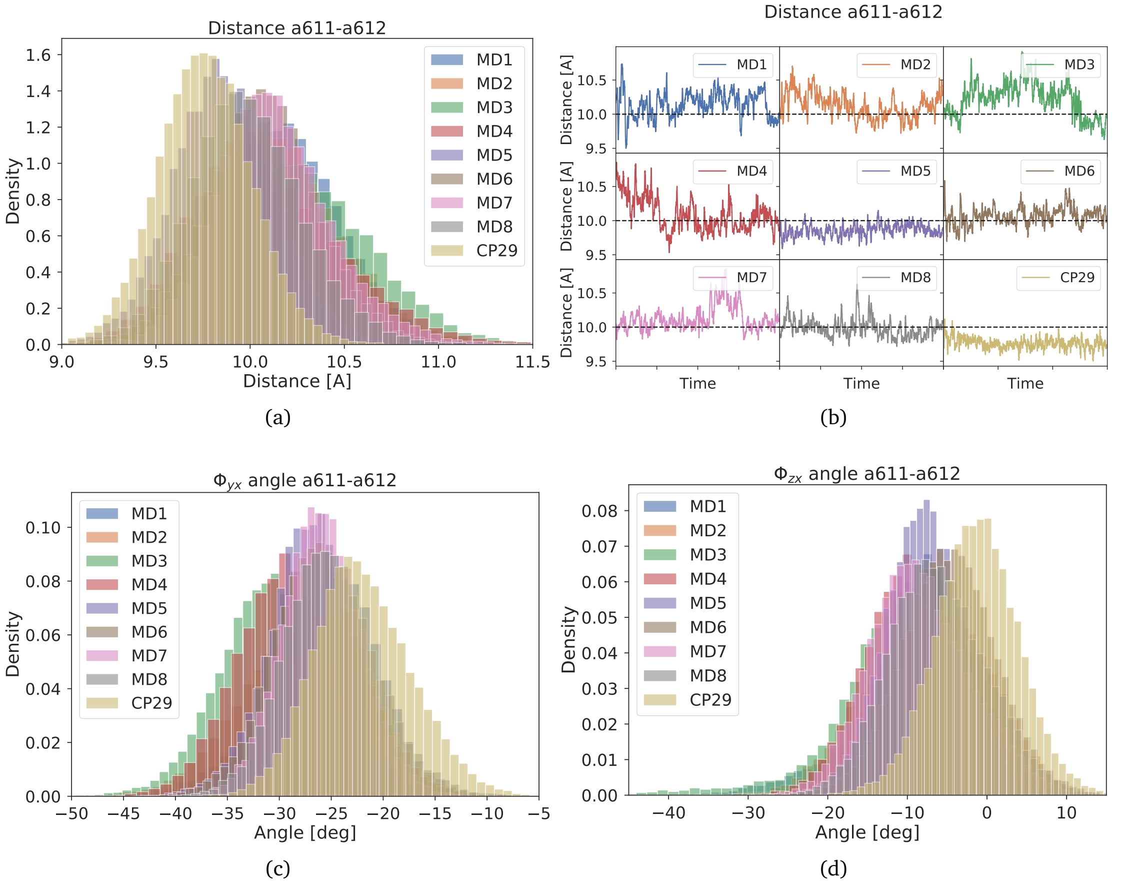


Figure S7: a-b) Mutual distance between central Mg site of the chlorophyll a611 and a612 for the individual replicas (MD) of the LHCSR1 and CP29. For the time evolution, we plotted running average over 100 frames c) Mutual rotation of the chlorophylls in the plane of the chlorophyll conjugated ring. d) Mutual rotation of the chlorophylls in the plane perpendicular to the conjugated ring.

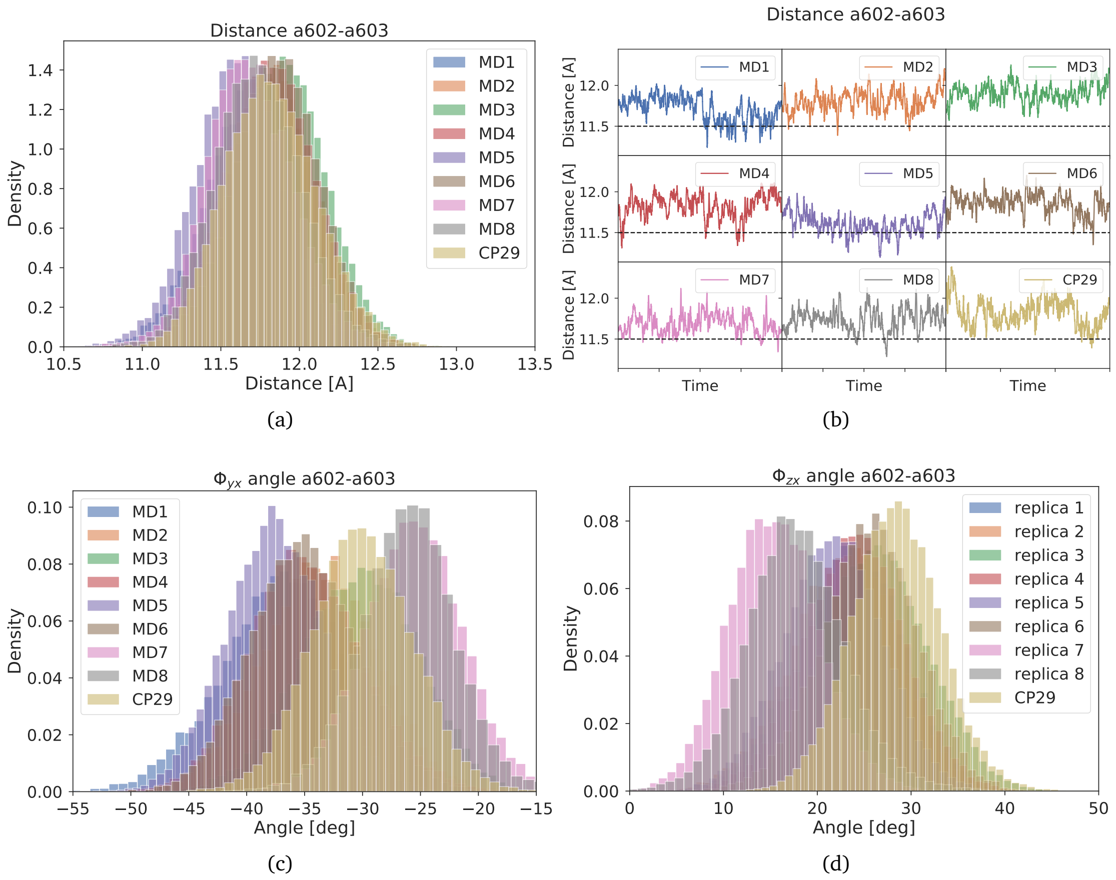


Figure S8: a-b) Mutual distance between central Mg site of the chlorophyll a602 and a603 for the individual replicas (MD) of the LHCSR1 and CP29. For the time evolution, we plotted running average over 100 frames c) Mutual rotation of the chlorophylls in the plane of the chlorophyll conjugated ring. d) Mutual rotation of the chlorophylls in the plane perpendicular to the conjugated ring.

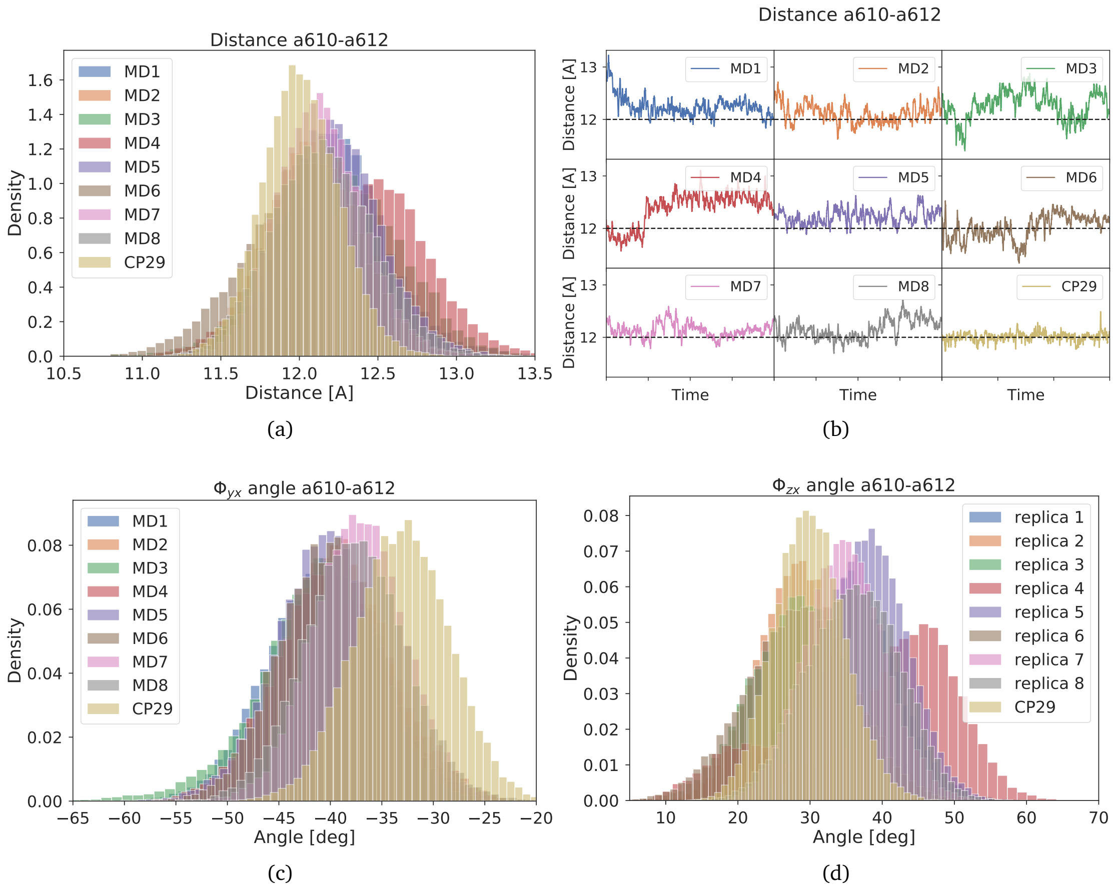


Figure S9: a-b) Mutual distance between central Mg site of the chlorophyll a610 and a612 for the individual replicas (MD) of the LHCSR1 and CP29. For the time evolution we plotted running average over 100 frames. c) Mutual rotation of the chlorophylls in the plane of the chlorophyll conjugated ring. d) Mutual rotation of the chlorophylls in the plane perpendicular to the conjugated ring.

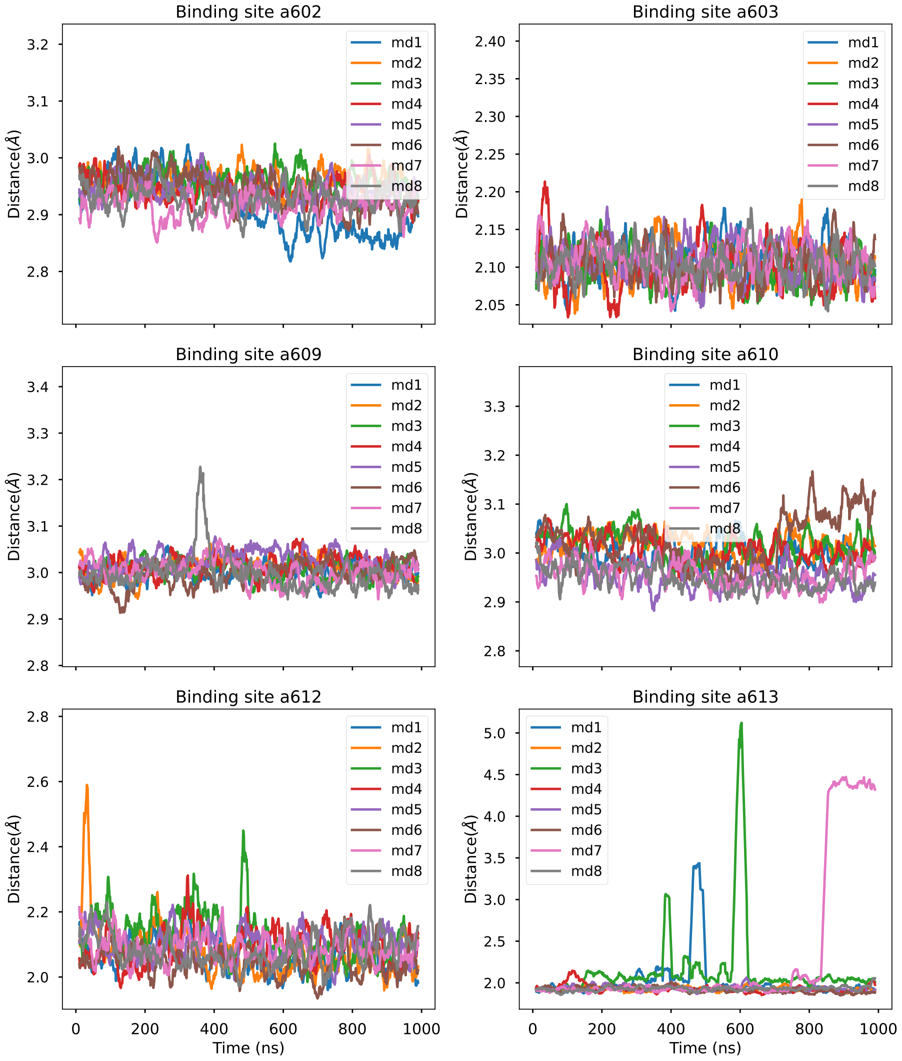


Figure S10: Distances from the Mg to the binding site atom for the Chls coordinated by a protein residue, along the eight MD replicas.

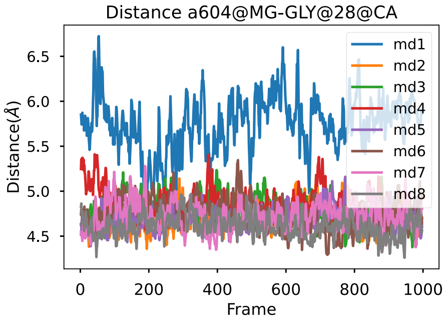


Figure S11: Distance from the Mg site of Chl a604 to the alpha carbon of the closest residue (Gly28) along the eight MD replicas.

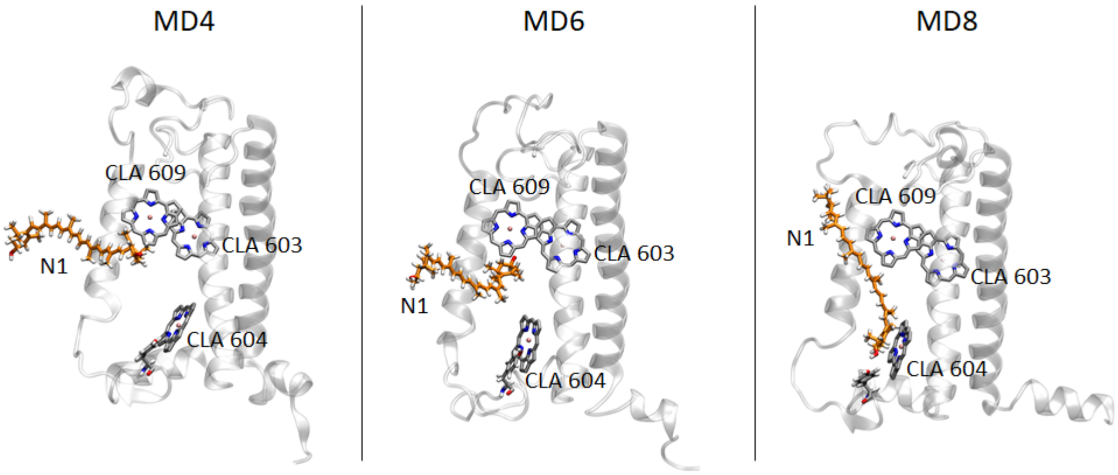


Figure S12: Representative configurations showing the position of N1-Lut in MD4, MD6, and MD8. Chlorophylls a603 a064 and a609 are also shown, together with the binding Tyr.

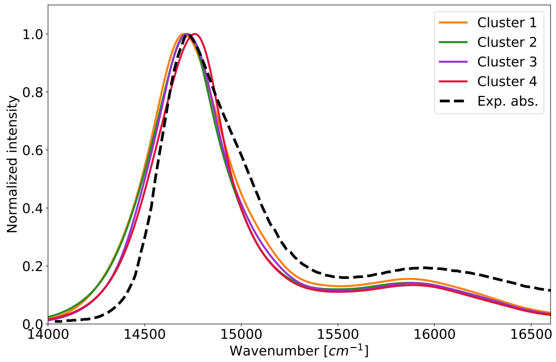


(a)

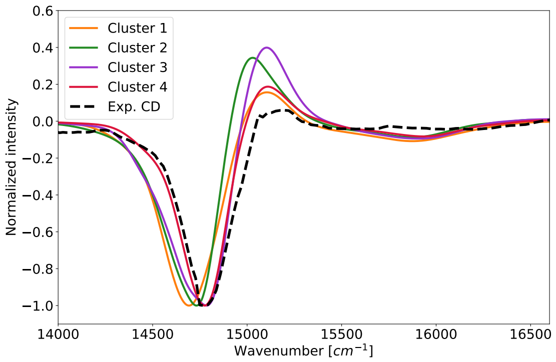


(b)

Figure S13: a) Absorption and b) CD spectra at 283K for the individual clusters of the LHCSR1. Best agreement with the experimental spectra Ref 5 is obtained for the cluster 4 which is than used in the main text for all the comparison with the CP29. The CD spectra are normalized to the main negative peak of the cluster 4 of the LHCSR1 system.

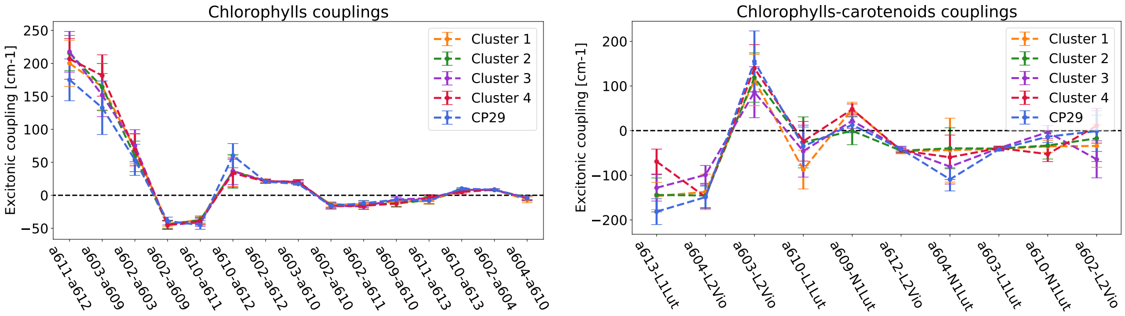


(a) (b)

Figure S14: Exciton couplings between individual pigments of the LHCSR1 compared with the couplings between corresponding pigments in the CP29. The couplings are ordered from the largest ones to the lowest (based on the cluster 1 of the LHCSR1). a) Exciton couplings between chlorophylls. b) Exciton couplings between chlorophylls and carotenoids.

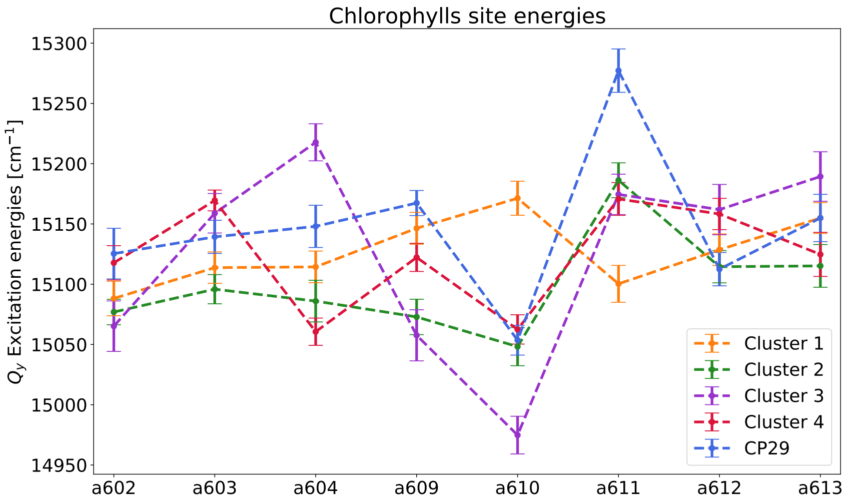


(a)

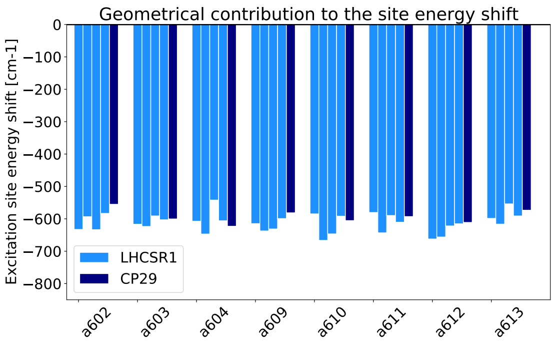


(b)

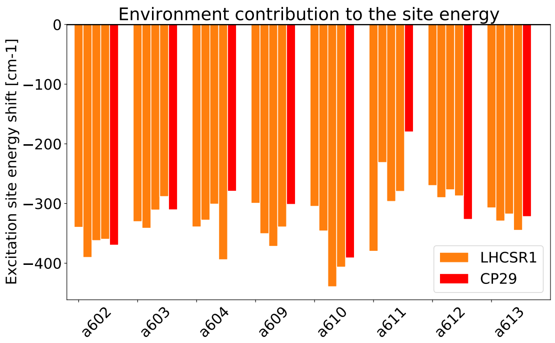


(c)

Figure S15: a) TD-DFT M06-2X site energies for the individual clusters of the LHCSR1 compared with the ones for the CP29. The error bars correspond to the standard error of the mean value. The site energy is decomposed into the b) geometrical and c) environmental contribution. The geometrical contribution is defined as the difference of the vacuum excitation energy from the optimized pigment in vacuum. The environmental contribution is the difference between vacuum and MMPol excitation energies, where the protein and solvent environment are included. All the presented site energies are after removing the bond-length fluctuations as described in the Supplementary Methods.

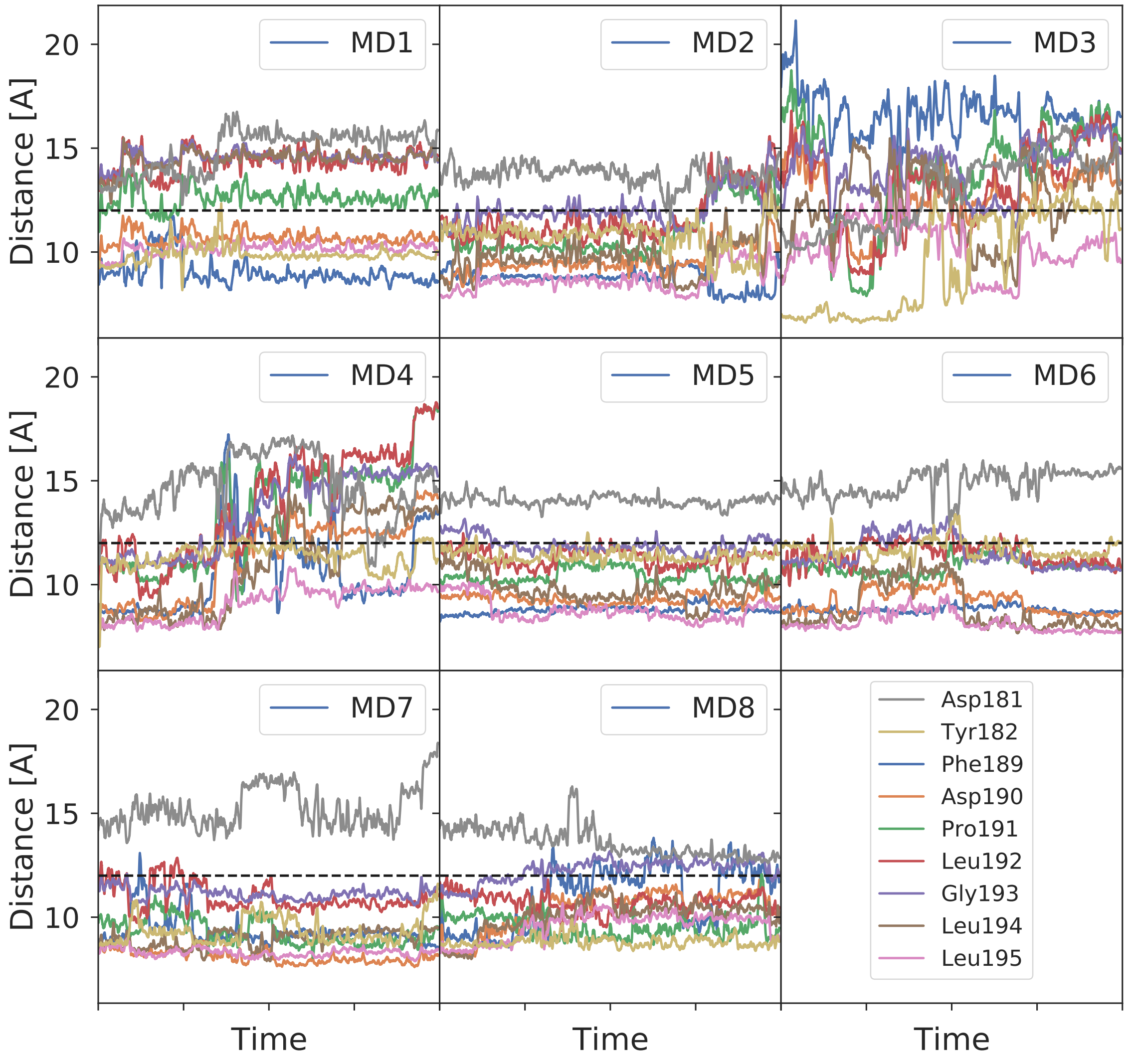


Figure S16: Mutual distance between chlorophyll a610 and selected residues from the stromal loop for each replica of the LHCSR1.

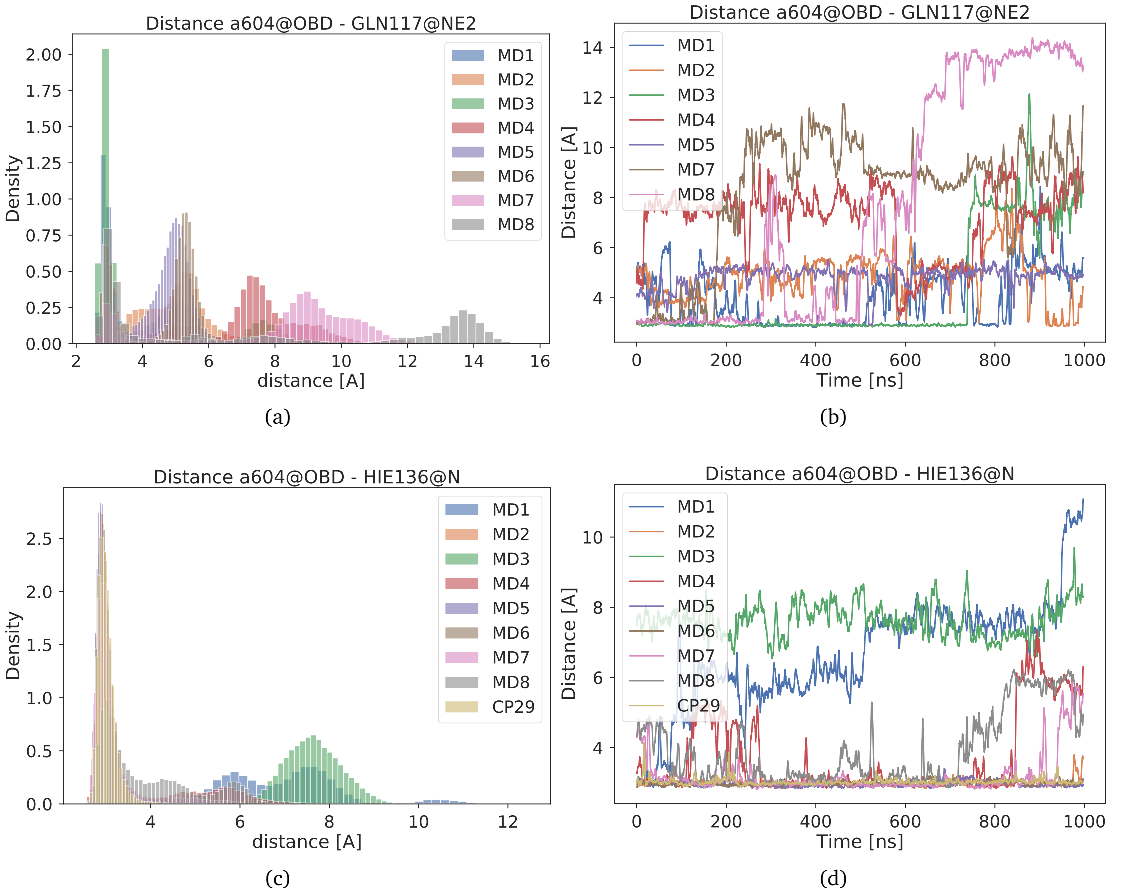


Figure S17: Mutual distance between chlorophyll a604 OBD oxygen which is part of the $\pi$-conjugated system and hydrogen bonding residues a-b) Gln117 and c-d) His136 for the LHCSR1 and corresponding residue for the CP29.

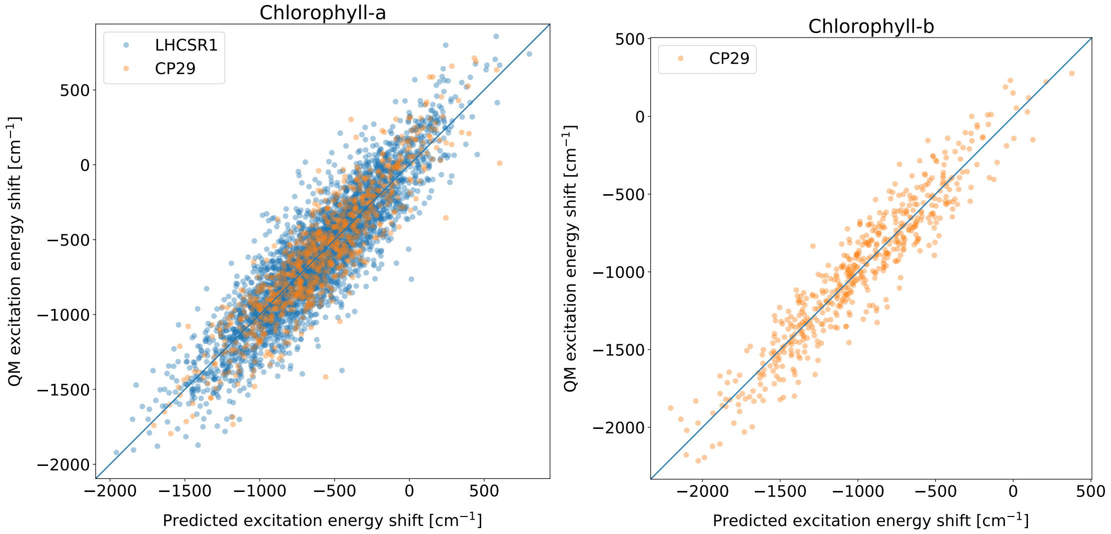


(a) (b)

Figure S18: Comparison of the fitted (predicted) vacuum excitation energy with the full QM one. The excitation energy is given as a difference with respect to the optimized structure of the pigment. For the fitting only bond-lengths for the pi-conjugation chain were used. The fitted vacuum excitation energies correspond very well to the QM ones. The difference between the fitted and the full QM excitation energy is interpreted as an angular (distortion) contribution to the vacuum site energy.

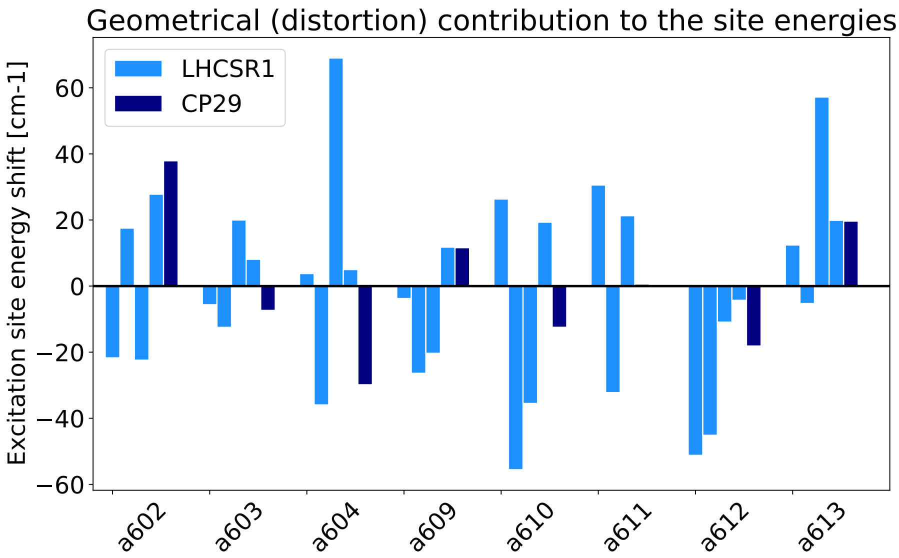


Figure S19: The averaged distortion contribution to the excitation energy for the LHCSR1 clusters and the CP29. The distortion contribution was computed as the difference of the predicted excitation energy and the full QM vacuum excitation energy, which was then averaged over the individual clusters.

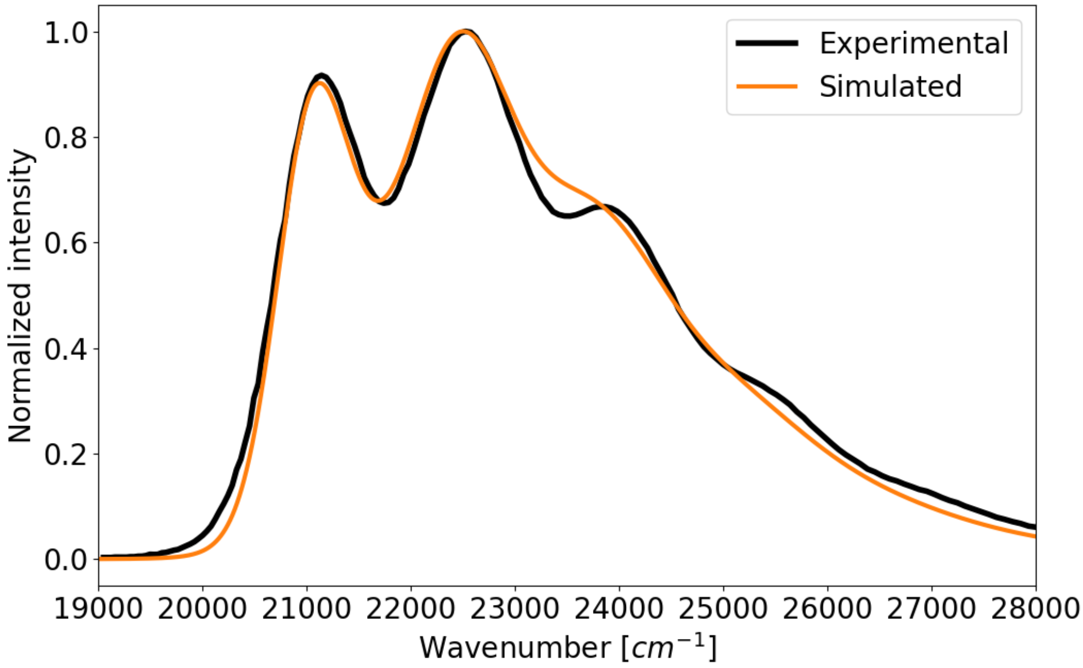


Figure S20: Comparison of the simulated and experimental[6] spectra of the Lutein in n-hexane which have the optical permitivity close to the one in the protein.

### Supplementary Tables

| | exp. peak position [$cm^{-1}$] | m06-2X [$cm^{-1}$] |
|---|---|---|
| Chl-a | 15105 | 17085 |
| Lutein | 21120 | 20261 |
| Violaxanthin | 21277 | 20559 |

Table S1: Comparison of the QM and experimental excitation energies for the LHCSR1 pigments in n-hexane. The TD-DFT energy gap between Chl-a and Cars computed with M06-2X functional is underestimated by $\sim 3000cm^{-1}$. Therefore the Cars are for the absorption spectra calculation all consistently shifted by the $3000cm^{-1}$ to get the correct experimental energy gap between Chl-a and Cars. This is needed to correctly evaluate the effect of the carotenoids on the optical spectra of both LHCSR1 and CP29.

| | a602 | a603 | a604 | a609 | a610 | a611 | a612 | a613 | L1-Lut | L2-Vio | N1-Lut |
|---|---|---|---|---|---|---|---|---|---|---|---|
| a602 | 15118 | 69 | 9 | -45 | -17 | -16 | 21 | -4 | 16 | 11 | -2 |
| a603 | 69 | 15170 | -3 | 182 | 21 | 1 | -3 | 6 | -39 | 140 | -17 |
| a604 | 9 | -3 | 15061 | -6 | -4 | -4 | 4 | 3 | 14 | -149 | -60 |
| a609 | -45 | 182 | -6 | 15122 | -13 | 5 | 0 | -3 | 17 | -3 | 48 |
| a610 | -17 | 21 | -4 | -13 | 15062 | -39 | 33 | 5 | -24 | 14 | -53 |
| a611 | -16 | 1 | -4 | 5 | -39 | 15171 | 207 | -3 | 13 | 13 | 9 |
| a612 | 21 | -3 | 4 | 0 | 33 | 207 | 15158 | 0 | 21 | -44 | -33 |
| a613 | -4 | 6 | 3 | -3 | 5 | -3 | 0 | 15125 | -69 | 18 | -5 |
| L1-Lut | 16 | -39 | 14 | 17 | -24 | 13 | 21 | -69 | 20202 | -62 | -27 |
| L2-Vio | 11 | 140 | -149 | -3 | 14 | 13 | -44 | 18 | -62 | 20180 | 106 |
| N1-Lut | -2 | -17 | -60 | 48 | -53 | 9 | -33 | -5 | -27 | 106 | 19621 |

Table S2: Exciton Hamiltonian of the LHCSR1 calculated within QM/MMpol approach at M062x/6-31G(d) level. All the values have been obtained as averages on the configurations extracted from MD and cluster 4, which represents the most equilibrated structure. The site excitation energies were shifted by -1381 $cm^{-1}$ and carotenoids by +1583 $cm^{-1}$. All values are in $cm^{-1}$

| | a601 | a602 | a603 | a604 | b606 | b607 | b608 | a609 | a610 | a611 | a612 | a613 | L1-Lut | L2-Vio | N1-Nex |
|---|---|---|---|---|---|---|---|---|---|---|---|---|---|---|---|
| a601 | 15146 | 13 | -5 | 0 | 0 | 0 | 0 | 0 | 1 | 30 | -6 | 1 | 3 | -36 | 0 |
| a602 | 13 | 15125 | 54 | 8 | 7 | 8 | -8 | -39 | -16 | -12 | 21 | -4 | 9 | -1 | -10 |
| a603 | -5 | 54 | 15139 | -4 | -8 | 4 | 3 | 133 | 17 | 0 | -3 | 1 | -42 | 155 | 0 |
| a604 | 0 | 8 | -4 | 15148 | 123 | 32 | -4 | -6 | -4 | -3 | 1 | 1 | 8 | -149 | -109 |
| b606 | 0 | 7 | -8 | 123 | 15402 | 35 | -4 | 1 | -3 | 0 | 2 | 1 | 0 | -83 | -104 |
| b607 | 0 | 8 | 4 | 32 | 35 | 15445 | -5 | -13 | 0 | 0 | 2 | 3 | -5 | -98 | -15 |
| b608 | 0 | -8 | 3 | -4 | -4 | -5 | 15420 | 46 | 79 | 7 | -1 | 0 | -8 | -9 | 36 |
| a609 | 0 | -39 | 133 | -6 | 1 | -13 | 46 | 15167 | -7 | 4 | -1 | -4 | 11 | 14 | 13 |
| a610 | 1 | -16 | 17 | -4 | -3 | 0 | 79 | -7 | 15054 | -45 | 60 | 10 | -35 | 15 | -13 |
| a611 | 30 | -12 | 0 | -3 | 0 | 0 | 7 | 4 | -45 | 15277 | 175 | -8 | 4 | 16 | 8 |
| a612 | -6 | 21 | -3 | 1 | 2 | 2 | -1 | -1 | 60 | 175 | 15112 | 1 | 54 | -41 | -18 |
| a613 | 1 | -4 | 1 | 1 | 1 | 3 | 0 | -4 | 10 | -8 | 1 | 15155 | -181 | 4 | -4 |
| L1-Lut | 3 | 9 | -42 | 8 | 0 | -5 | -8 | 11 | -35 | 4 | 54 | -181 | 20167 | -74 | -12 |
| L2-Vio | -36 | -1 | 155 | -149 | -83 | -98 | -9 | 14 | 15 | 16 | -41 | 4 | -74 | 20323 | 101 |
| N1-Nex | 0 | -10 | 0 | -109 | -104 | -15 | 36 | 13 | -13 | 8 | -18 | -4 | -12 | 101 | 20737 |

Table S3: Exciton Hamiltonian of the CP29 calculated within QM/MMpol approach at M062x/6-31G(d) level. All the values have been obtained as averages on the configurations extracted from MD in Ref 7. The site excitation energies of the chlorophylls were shifted by -1381 $cm^{-1}$ and carotenoids by +1583 $cm^{-1}$. All values are in $cm^{-1}$.

| exciton n. | LHCSR1 | CP29 |
|---|---|---|
| 1 | 14910 | 14929 |
| 2 | 14947 | 14992 |
| 3 | 15054 | 15088 |
| 4 | 15085 | 15095 |
| 5 | 15125 | 15146 |
| 6 | 15153 | 15151 |
| 7 | 15329 | 15161 |
| 8 | 15371 | 15277 |
| 9 | | 15390 |
| 10 | | 15404 |
| 11 | | 15445 |
| 12 | | 15489 |
| 13 | 19603 | 20143 |
| 14 | 20139 | 20340 |
| 15 | 20274 | 20770 |

Table S4: TD-DFT M06-2X/MMPol exciton energies (in $cm^{-1}$) determined from MD structures for the LHCSR1 cluster 4 and CP29. The exciton energies were computed from the averaged Hamiltonian and therefore represent only approximate exciton energies to the ones used for the optical spectra calculation, which were computed separately for the Hamiltonian of each cluster and frame.
